# Functional dynamics of de-afferented early visual cortex in glaucoma

**DOI:** 10.1101/2020.09.16.300012

**Authors:** Gokulraj T. Prabhakaran, Khaldoon O. Al-Nosairy, Claus Tempelmann, Markus Wagner, Hagen Thieme, Michael B. Hoffmann

## Abstract

fMRI studies in macular degeneration (MD) and retinitis pigmentosa (RP) demonstrated that responses in the lesion projection zones (LPZ) of V1 are task related, indicating significant limits of bottom-up visual system plasticity in MD and RP. In advanced glaucoma (GL), a prevalent eye disease and leading cause of blindness, the scope of visual system plasticity is currently unknown. We performed 3T fMRI in patients with extensive visual field defects due to GL (n=5), RP (n=2) and healthy controls (n=7; with simulated defects). Participants viewed contrast patterns drifting in 8 directions alternating with uniform gray and performed 3 tasks: (1) passive viewing (PV), (2) one-back task (OBT) and (3) fixation-dot task (FDT). During PV, they passively viewed the stimulus with central fixation, during OBT they reported the succession of the same two motion directions, and during FDT a change in the fixation color. In GL, LPZ responses of the early visual cortex (V1, V2 and V3) shifted from negative during PV to positive for OBT [p (corrected): V1(0.006); V2(0.04); V3(0.008)], while they were negative in the controls’ simulated LPZ for all stimulation conditions. For RP a similar pattern as for GL was observed. Consequently, activity in the de-afferented visual cortex in glaucoma is, similar to MD and RP, task-related. In conclusion, the lack of bottom-up plasticity appears to be a general feature of the human visual system. These insights are of importance for the development of treatment and rehabilitation schemes in glaucoma.

**Highlights:**

1. Functional dynamics of early visual cortex LPZ depend on task demands in glaucoma
2. Brain activity in deprived visual cortex suggests absence of large-scale remapping
3. Limited scope of bottom-up plasticity is a general feature of human visual system
4. Visual system stability and plasticity is of relevance for therapeutic advances

## 1. Introduction

Glaucoma, a progressive degeneration of retinal ganglion cells (RGCs), results in an irreversible loss of vision, eventually leading to blindness (Jonas et al., 2017). Worldwide it is the second most prevalent cause of acquired blindness, next to cataract (Quigley and Broman, 2006). Most of the conventional therapeutic strategies, by medication or surgeries, are directed towards the management or control of the major risk factor in glaucoma, increased intra-ocular pressure (IOP). In fact, these interventions are known to reduce the progression rate of the disease (Jonas et al., 2017). However, the beneficial effects of recent advances in early detection and progression-delaying treatment of glaucoma are counteracted by increased life expectancy. As a consequence, a substantial proportion of glaucoma patients will become severely visually impaired and eventually bilaterally blind during their lifetime (Kapetanakis et al., 2016; Quigley and Broman, 2006). This makes it important for current research initiatives, not to only focus on better disease management tools, but also to further our understanding of the management of visual impairment in advanced glaucoma with the ultimate goal to explore effective avenues for the restoration of visual input.

In addition to the retinal damage caused by glaucoma, the concomitant deprivation of visual input from the retina to the cortex has been shown to result in structural and functional changes at the cortical level, in particular in primary (V1) and extra-striate (V2 & V3) visual cortex (Boucard et al., 2016; Dai et al., 2013; Duncan et al., 2007; Frezzotti et al., 2014; Wang et al., 2016; Zhou et al., 2017). fMRI-based retinotopic mapping has previously demonstrated the interplay of plasticity and stability in congenital visual pathway abnormalities and retinal diseases (Ahmadi et al., 2020, 2019; Hoffmann et al., 2012; Hoffmann and Dumoulin, 2015). In contrast, primary visual cortex reorganization is much more limited in patients with acquired visual field defects as studied in detail for in macular degeneration (Baker et al., 2008; Baseler et al., 2011; Dilks et al., 2009; Liu et al., 2010; Masuda et al., 2008; Plank et al., 2013). Although glaucoma is a prevalent disease, the scope of cortical plasticity in the deafferented portions of the early visual cortex in advanced glaucoma has only received little attention. Zhou et al. reported an enlarged para-foveal representation in the visual cortex of glaucoma patients (Zhou et al., 2017), but recent investigations of simulated peripheral response dropouts in controls (Prabhakaran et al., 2020) suggest that such effects do not necessarily reflect veridical long-term cortical reorganization. This mirrors the views based on previous reports analyzing the limited nature of visual cortex plasticity for foveal de-afferentation (Barton and Brewer, 2015; Baseler et al., 2011; Haak et al., 2012). Accordingly, reductions of amplitude and extent of fMRI BOLD responses in the visual cortex in glaucoma patients (Borges et al., 2015; Duncan et al., 2007; Murphy et al., 2016; Song et al., 2012) and thinning of gray matter in the de-afferented visual cortex were reported (Boucard et al., 2016; Yu et al., 2015, 2014), which also suggests the absence of large-scale reorganization post visual field loss. However, in the context of vision restoration and rehabilitation strategies, investigations of the lesion projection zones (LPZ) in the visual cortex representing the retinal lesions are key to understanding the reality of adult visual cortex reorganization capabilities. Consequently, investigations are needed to look at the scope of cortical responses in the LPZ and modulations in these responses, if any, in relation to visual stimulation and visual tasks. In fact, for patients with non-glaucomatous retinal disorders, cortical activations in the LPZ were reported. Remarkably they appear to depend on the presence of a visual task to be performed on the presented stimuli. In a series of case observations Masuda et al., demonstrated these task-dependent V1-responses in patients with macular degeneration (MD) and retinitis pigmentosa (RP), i.e. for central (Masuda et al., 2008) and peripheral visual field defects (Masuda et al., 2010), respectively. They explained these signals in the absence of long-term plasticity, e.g. as side effects of task-related feed-back from higher visual areas. Although highly relevant for the management of advanced glaucoma, such insights into the responsiveness of the LPZ in the early visual cortex are currently completely missing for the entity of glaucoma patients.

In the present study, we pursued a comparative approach to characterize such abnormal cortical responses in the de-afferented early visual cortex in glaucoma. We compared the task-dependence of cortical responses in the LPZ in patients with glaucoma to those with advanced RP and controls with simulated visual field defects. In fact, we found similar response signatures and task-dependencies in RP and glaucoma, which are analogous to those reported previously in MD, but absent in controls. Consequently, the lack of relevant bottom-up plasticity appears to be a general feature of the human visual system.

## 2. Methods

### 2.1 Participants

Five patients with extensive visual field (VF) deficits due to advanced open-angle glaucoma (age: 51 to 78; 3 males and 2 females), two female patients with advanced changes due to retinitis pigmentosa (age: 46 & 56) and seven age-matched visually healthy individuals with normal vision (best-corrected decimal visual acuity ≥ 1.0 (Bach, 1996)) participated in the study. Table 1 shows the detailed demographics of the participants. Written informed consents and data usage agreements were signed by all participants. Within the limits of privacy issue of clinical data, the data will be made available via a repository. The study was conducted in adherence to the tenets of the Declaration of Helsinki and was approved by the ethics committee of the University of Magdeburg.

**Table 1.** Participant demographics and clinical characteristics. All measures are for the stimulated eye in fMRI; Visual field mean deviation (MD) as measured with standard automated perimetry (SAP); Fixation stability within 2° visual angle of the fixation target as measured with microperimetry; na – not available.

| Participant | Age | Gender | Diagnosis | Stimulated eye - fMRI | Visual acuity | Visual field (MD - dB) | Fixation stability (2°) |
| --- | --- | --- | --- | --- | --- | --- | --- |
| GL1 | 68 | female | glaucoma | left | 1.0 | -13.9 | 100% |
| GL2 | 78 | male | glaucoma | left | 0.05 | -30.7 | 74% |
| GL3 | 70 | male | glaucoma | left | 0.4 | -25.8 | 96% |
| GL4 | 51 | female | glaucoma | right | 0.8 | -32.5 | na |
| GL5 | 72 | male | glaucoma | left | 0.8 | -30.7 | 99% |
| RP1 | 46 | female | retinitis pigmentosa | right | 0.05 | -27.3 | 5% |
| RP2 | 56 | female | retinitis pigmentosa | left | 0.6 | -23.5 | 91% |
| C1 | 64 | male | control | left | 1.6 | 1.0 | na |
| C2 | 79 | female | control | left | 1.3 | 0.0 | na |
| C3 | 56 | male | control | left | 1.3 | 0.0 | na |
| C4 | 73 | male | control | left | 1.0 | -0.9 | na |
| C5 | 75 | male | control | left | 1.3 | 0.0 | na |
| C6 | 71 | female | control | right | 1.0 | -1.4 | na |
| C7 | 63 | female | control | left | 1.0 | na | na |

### 2.2 Visual field testing and fixation stability

Standard automated perimetry was performed using 24-2 Swedish Interactive Threshold Algorithm protocol (SITA-Fast; Goldmann size III white-on-white stimuli; Humphrey Field Analyzer 3; Carl Zeiss Meditec AG; Jena, Germany). For two patients, VF to the central 30° was tested using another perimeter (OCTOPUS^®^ Perimeter 101, Haag-Streit International, Switzerland; dG2; dynamic strategy; Goldmann size III). For the patient cohort (except 1; GL4), fixation stability was determined with a fundus-controlled microperimeter (MP-1 microperimeter, Nidek, Padova, Italy). Deviations from central fixation were tracked (at 25 Hz sampling frequency) and the proportion of fixations falling within central 2° of retina was reported using built-in MP1 analysis (Table 1).

### 2.3 Visual stimulation for fMRI

#### Stimulus conditions and rationale

Three different tasks were performed independently in separate runs within the same session: (1) one-back task (OBT), (2) passive viewing (PV) and (3) fixation-dot task (FDT). The underlying rationale was to dissociate top-down modulations and bottom-up input to the visual cortex by applying visual stimulation with a moving pattern with (OBT) and without a stimulus related task (PV and FDT). Specifically, during (1) OBT, the participants were instructed to report a repetition of same drifting directions of the pattern in two consecutive trials using a button press. They were required to fixate on the central dot while performing the task. One-back repetition trials were at least 15% of the total number of trials and were randomized. All the participants were able to perform the stimulus-locked task without much difficulty. (2) During PV, they passively viewed the stimulus fixating on the central dot, i.e. they did not perform any task, and they were explicitly instructed not to do the OBT during the PV. During (3) FDT, a fixation dot task (not locked to the stimulus, i.e. running during both on- and off-blocks) was added to ensure OBT was not performed. For FDT the participants responded via button press when the color of the fixation dot changed. In all controls and most of the patients, the switch-colors used were black and white; however, in some patients different colors were used depending on the ability of participants to notice the change. The color change occurred throughout the cycle i.e. during both the stimulus presentation and the mean luminance gray. The spatial and temporal properties of the stimuli were kept consistent for all the three conditions. The underlying aim of including FDT was to compare PV and FDT in order to validate that the condition PV was sufficient to achieve passive viewing with respect to the stimulus as for FDT. In that case similar activation patterns are expected for PV and FDT.

#### Visual stimulation

Psychtoolbox-3 (Brainard, 1997; Pelli, 1997) was used to program the visual stimuli in MATLAB (Mathworks, Natick, Massachusetts, USA). The stimulus employed comprised high-contrast patterns drifting in eight different directions that were projected to a screen at the rear end of the magnet bore, with a resolution of 1920 x 1080 pixels. Participants viewed the stimulus monocularly via their better eye at a distance of 35 cm via an angled mirror. This resulted in an effective stimulus size subtending approximately 24° and 14° radius in the horizontal and the vertical directions, respectively. All the patients viewed the stimulus projected on the entire screen, whereas, in the controls, we simulated an artificial peripheral scotoma by exposing only the central 7° of the stimulus through a circular aperture. The temporal sequence of each run followed a block design with 10 cycles of 12 s motion block (stimulus presentation) alternating with 12 s of mean luminance gray (24 s per cycle). Within each motion block, the direction of the contrast pattern was randomly changed every second (i.e. 12 trials per block; Figure 1). In each 1 s trial, the stimulus was presented for 750 ms followed by a 250 ms mean luminance gray. Participants were instructed to maintain fixation on a centrally located fixation dot.

**Fig 1.**
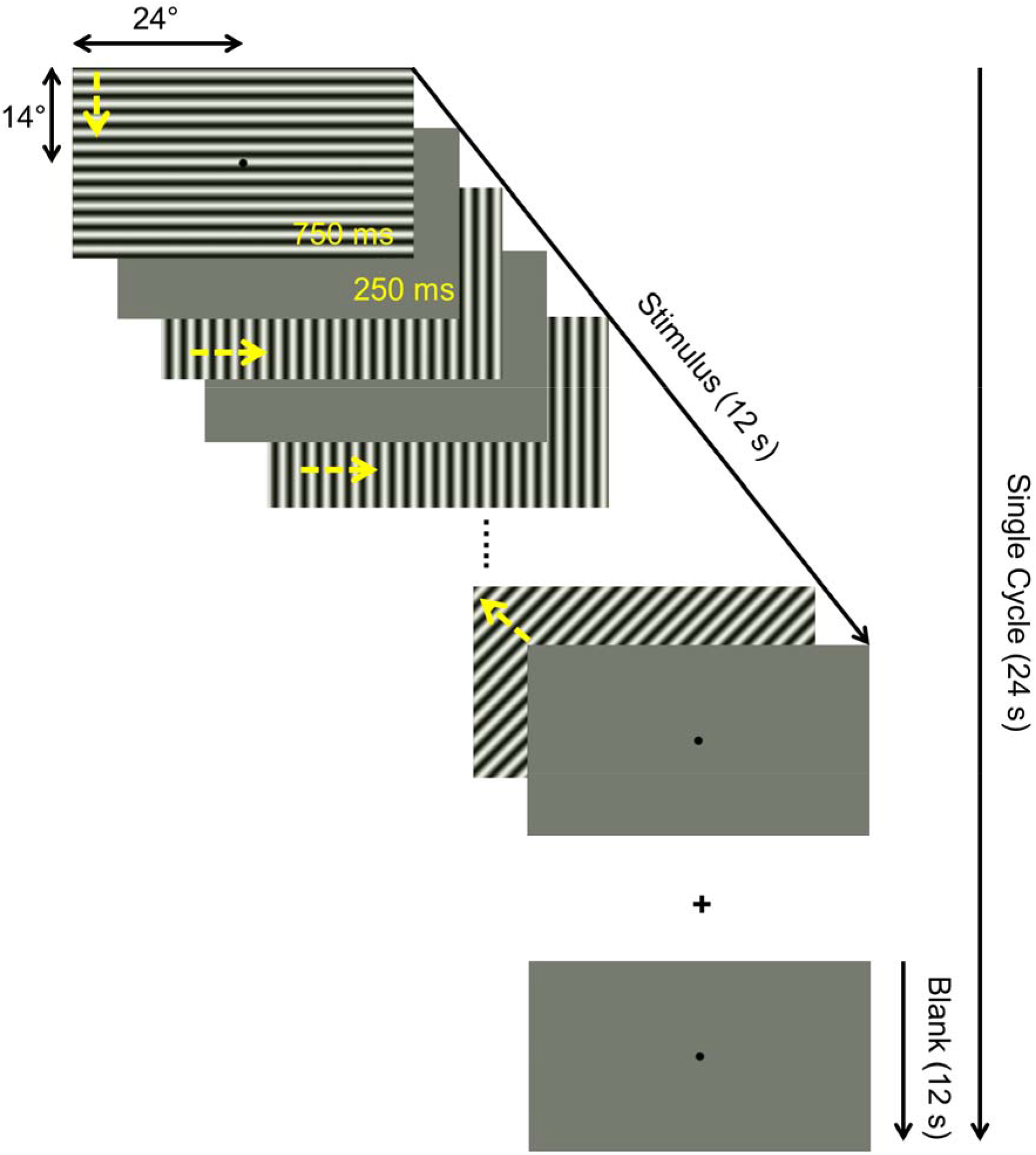
Illustration of temporal sequence of a single stimulus cycle. Each 24 s cycle comprised a 12 s motion block (drifting patterns) block and a 12 s mean blank block (mean luminance grey). A total of 10 cycles were presented per fMRI run, resulting in a total duration of 240 s. The motion block consisted of 12 one-second trials comprising 750 ms motion stimulus (drifting in one of 8 different directions, as indicated by yellow arrows added for visualization) and 250 ms mean luminance grey. Identical stimuli were presented for three different task conditions in separate fMRI runs: (i) passive viewing (PV), (ii) one-back task (OBT), (iii) fixation-dot task (FDT), as detailed in Methods.

In addition, to delineate the visual areas, fMRI based population receptive field (pRF) mapping scans were obtained from each participant in a second session on a separate day. A checkerboard stimulus pattern (mean luminance: 109 cd/m^2^; contrast: 99%; check size: 1.57°) moving in eight different directions (2 horizontal, 2 vertical and 4 diagonal; Dumoulin and Wandell, 2008) was exposed through a bar aperture. The width of the bar subtended 1/4^th^ (3.45°) of the stimulus radius (13.8°). The spatial and temporal properties of the stimulus have been described in Prabhakaran et al. (Prabhakaran et al., 2020). The duration of each pRF mapping scan was 192 s and the scan was repeated 6 times for the patient cohort and 4 times for the controls. The participants responded to a fixation-dot color change via button press.

### 2.4 MRI acquisition

All MRI and fMRI data were collected on a 3 Tesla Siemens Prisma scanner (Erlangen, Germany). In order to allow for an unrestricted view of the entire projection screen, we used only the lower section of a 64-channel head coil, resulting in a 34-channel coil covering most of the brain. fMRI scans parallel to the AC-PC line were acquired using a T2*-weighted BOLD gradient-EPI sequence (TR | TE = 1500 | 30 ms & voxel size = 2.5^3^ mm^3^). A total of 160 fMRI time series images (volumes) were obtained for each run after removal of the first 8 volumes by the scanner itself to allow for steady magnetization. The fMRI parameters were the same for the pRF mapping data, except for the number of volumes (136). One high-resolution whole brain anatomical T1-weighted scan (MPRAGE, 1 mm isotropic voxels, TR | TI | TE = 2500 | 1100 | 2.82 ms) was collected for each participant to allow for cortical visualization of fMRI responses. An inversion recovery EPI sequence (TR | TI | TE = 4000 | 1100 | 23 ms) with spatial coverage (FOV) and resolution identical to the T2* EPI was obtained to aid in the alignment of structural and functional data.

### 2.5 Data preprocessing

Gray-white matter boundaries in the T1-weighted anatomical images were segmented using Freesurfer (https://surfer.nmr.mgh.harvard.edu/). ITK-gray (https://github.com/vistalab/itkgray) was used to manually inspect the Freesurfer segmentation and correct for possible segmentation errors. A 3-D rendering of the cortical surface was generated by reconstruction of the segmented boundaries (Wandell et al 2000). Within and between fMRI scans, head motion artefacts were corrected using AFNI (https://afni.nimh.nih.gov/). For each participant, motion-corrected fMRI time series of the repetitions of each stimulation condition (i.e. PV, OBT, FDT, and pRF mapping data) were averaged into separate groups with MATLAB-based Vistasoft tools (mrVista https://github.com/vistalab/vistasoft). The inversion recovery EPI was aligned spatially with the anatomical scan in two steps; first manually with rxAlign function in mrVista and then automatically using Kendrick Kay’s alignment toolbox (https://github.com/kendrickkay/alignvolumedata). The obtained alignment matrix was used to align the fMRI images with the anatomy.

### 2.6 Data analysis

#### Phase-specified coherence (Coherence_ps_)

We computed voxel-wise coherence at the fundamental stimulus frequency and phase corrected for hemodynamic delay (phase-specified coherence) to quantitatively investigate changes in the strength of the BOLD response for the different task conditions. We used the definition and formula to calculate the phase-specified coherence (coherence_*ps*_) that has been previously used (Masuda et al., 2008) and is also available as a function in the mrVista toolbox,

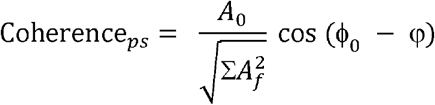

where A_0_ and ϕ_0_ are the signals amplitude and phase at the stimulation frequency, respectively, Af are the amplitudes of each Fourier component, and φ is the delay in the hemodynamic response (estimated for each participant from the positive fMRI responses). The phase at the stimulus frequency was estimated from the averaged fMRI time-series of all the task conditions in a small 5mm ROI drawn in the region of the cortex that had reliable positive BOLD response across all the conditions. Coherence_*ps*_ can take values between −1 and +1; voxels with positive measure reflect stimulus synchronized fMRI response modulation and negative measures reflect modulation to the mean luminance gray (no or negative stimulus related BOLD response).

#### Visual area delineation

We defined the borders of primary (V1) and extra-striate visual cortex (V2 & V3) for each participant using fMRI-based pRF-mapping data. Employing a 2D-Gaussian pRF model approach described previously (Dumoulin and Wandell, 2008; Prabhakaran et al., 2020), we estimated for each voxel their preferred position in the visual field (x and y in Cartesian coordinates). Eccentricity 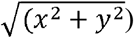 and polar angle 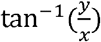 measures were derived from these position parameters. Polar angle estimates were projected onto an inflated cortical surface and the visual areas were delineated by following the phase reversals in the polar angle data (Sereno et al., 1995). Unlike controls, this delineation process was not straightforward in the patients because of retinal and subsequent cortical lesions. So in patients with incomplete cortical representation, we used the Benson atlas (Benson et al., 2014), which for every individual applies an anatomically defined template of retinotopy, to assist in our pRF-mapping based visual area definitions. The anterior extent of the visual areas was manually drawn based on the participants pRF mapping data and Benson atlas extracted eccentricity predictions (14° in the vertical meridian representation and 24° in the horizontal meridian representation) in correspondence to our stimulus size. Based on the coherence_*ps*_, measures from PV we divided each visual area into two ROIs; voxels with positive responses were classified as the normal projection zone (NPZ) and those with negative responses as the lesion projection zone (LPZ).

All the further region of interest (ROI) analyses were performed with custom written scripts in MATLAB and statistics in SPSS 26 (Statistical Package for the Social Sciences, IBM) and corrected for multiple comparison with the Holm-Bonferroni correction (Holm, 1979).

## 3. Results

In a comparative approach with visually healthy individuals and simulated peripheral scotoma, we investigated the scope of aberrant cortical responses in the de-afferented visual cortex of glaucoma patients with advanced VF deficits. As a validation and replication to the existing literature, we also examined such responses in a second reference group of two patients with RP.

A qualitative overview of the impact of task on the fMRI-responses in the visual cortex of a representative glaucoma participant is given in Figure 2, and for a control and an RP patient in Figure 3 and suppl. Figure S1, respectively. It is shown that in glaucoma, a specific task dependent signature is evident in the de-afferented cortex, i.e. the LPZ, while such task dependence is absent in the control with a simulated peripheral scotoma. While visual stimulation in the unstimulated LPZ induced negative BOLD responses for all conditions tested, the LPZ-BOLD-signature in glaucoma depended on task. Specifically, instead of the negative LPZ-BOLD responses for PV, positive BOLD responses were obtained in LPZ during the OBT in glaucoma. Notably, response patterns for PV and FDT did not differ. For the RP participant (suppl. Figure S1) a similar pattern of task-dependent responses in the LPZ as for the glaucoma patient was observed. In summary, the time series modulations in the glaucoma patient’s LPZ was comparable to the controls during PV and FDT, but not for the OBT. For the latter, the glaucoma patient’s LPZ BOLD response during stimulation was shifted from negative to positive, as for the RP patient. In addition, an increase in the amplitude of the NPZ BOLD response was also observed for the OBT in the glaucoma patient.

**Fig 2.**
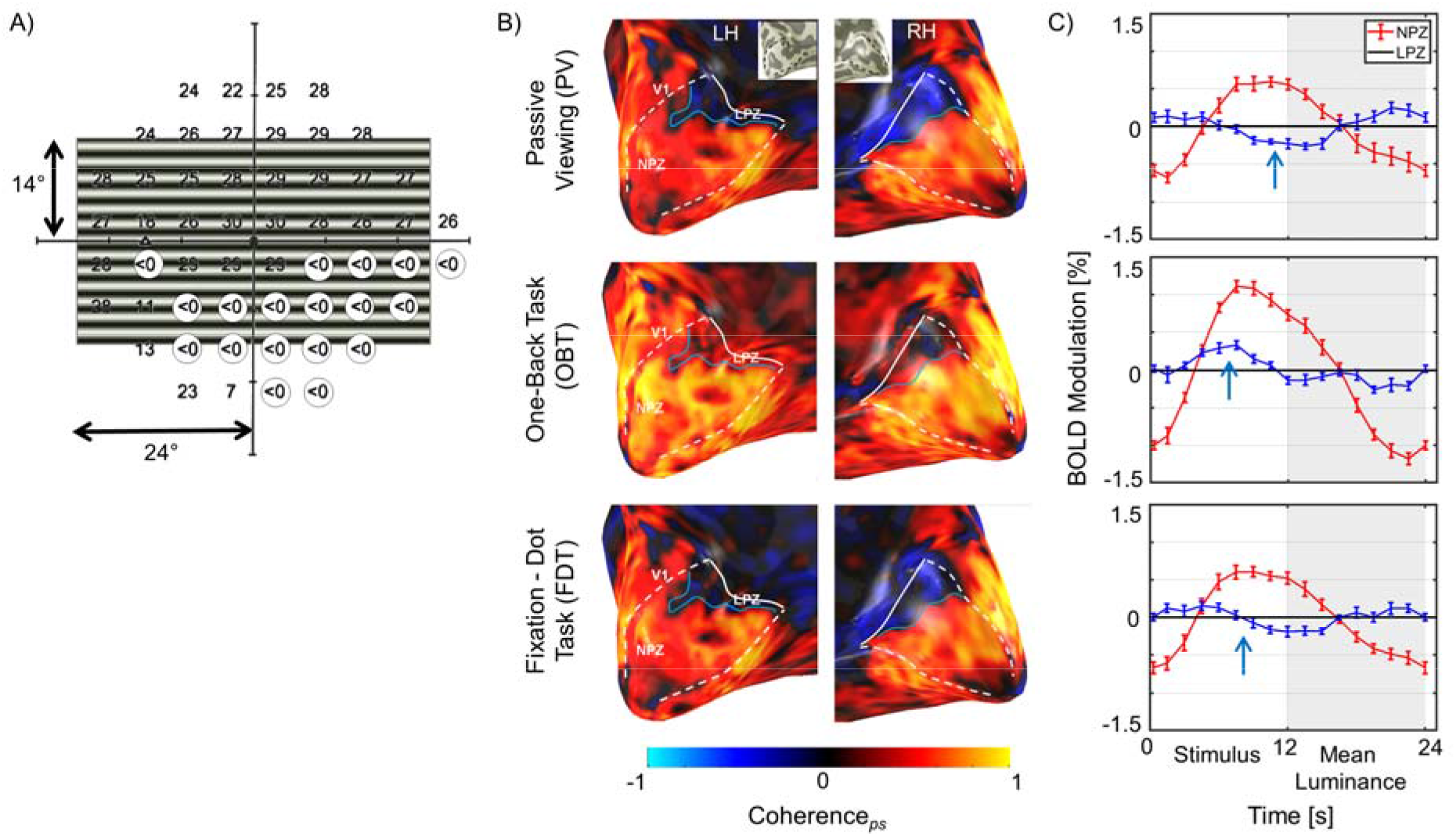
Visual field and fMRI-activations for glaucoma participant GL1 (left eye). **A)** Visual field defect. The visual field sensitivities for the left eye (stimulated in the fMRI experiment) as determined perimetrically are superimposed onto the stimulus layout (absolute scotomas are highlighted by white discs). **(B)** BOLD-activations (coherence_*ps*_) for the three task conditions projected onto the inflated occipital lobe as false-color overlays. V1 boundaries (white dashed) and the anterior extent (white solid line) were determined from the participants pRF mapping data informed by atlas mapping as detailed in Methods. NPZ – normal projection zone; LPZ – lesion projection zone. **(C)** Average single-cycle BOLD time series for the three conditions in the NPZ (red) and LPZ (blue) ROIs. White and grey zones indicate motion and blank blocks, respectively. The induced BOLD response is shifted due to the hemodynamic delay. Depending on tasks, LPZ responses are negative (PV) or positive (OBT) or reduced (FDT), as indicated by the arrows.

**Fig 3.**
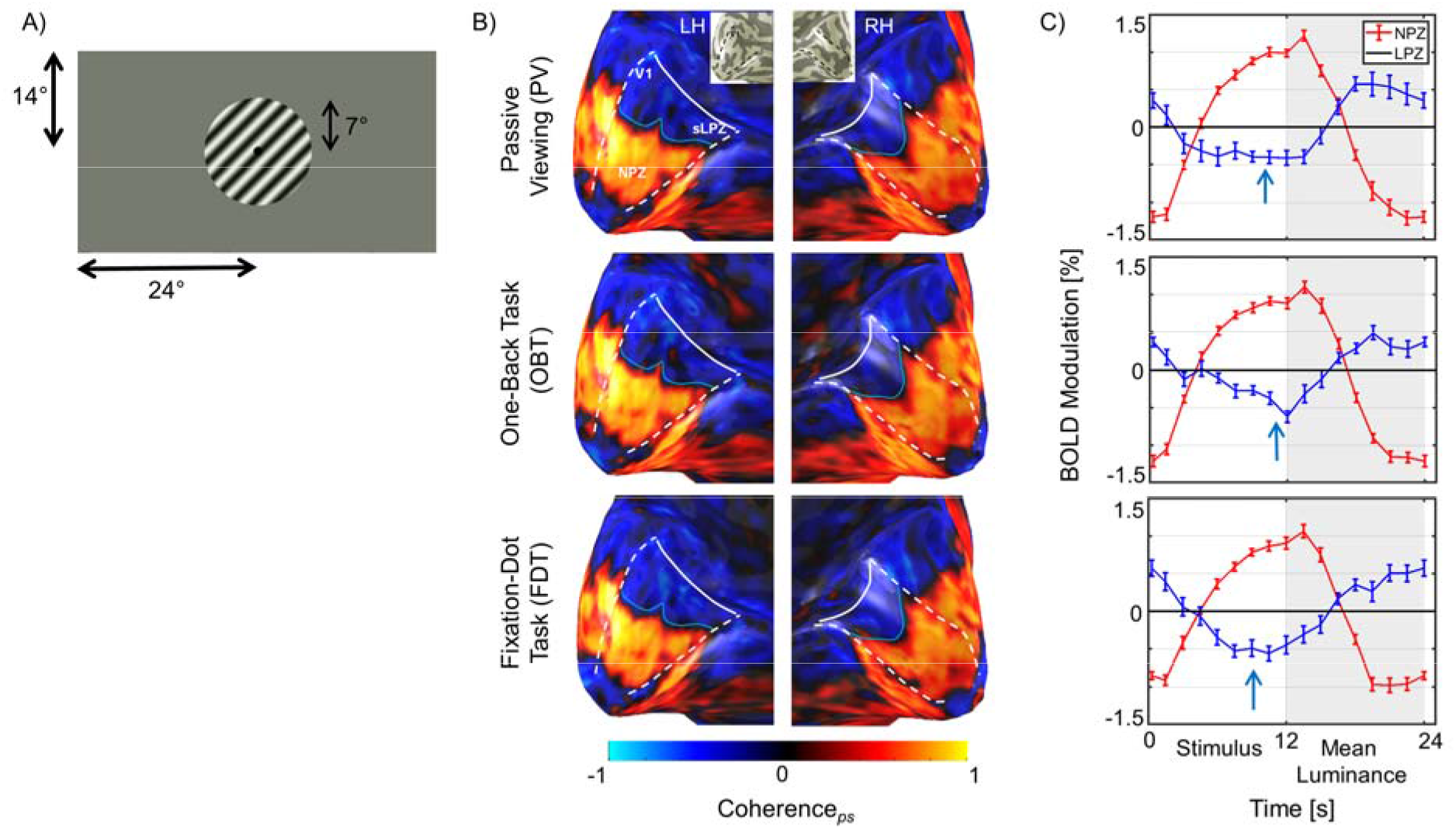
fMRI-activations for control C1 (right eye). **(A)** Illustration of stimulated visual field for the controls with a scotoma simulation by peripheral masking beyond 7°. **B)** BOLD-activations (coherence_*ps*_) as false-color overlays onto the inflated occipital lobe. **C)** Average single-cycle BOLD time series. Sign of stimulus induced LPZ responses are negative, irrespective of the task condition. Conventions as for Figure 2.

In a quantitative assessment of the group data the above task dependence of the LPZ-responses was further confirmed for glaucoma and also demonstrated for RP. For each participant, within V1, we quantified the mean phase-specified coherence of the voxels in the NPZ and LPZ for the three different conditions. In Figure 4A the group mean coherence_*ps*_, for controls (n=7), glaucoma (n=5), and RP (n=2) are depicted (see suppl. Figure S2 for subjective measures). Irrespective of participant group, we observed a strong positive coherence_*ps*_. in the NPZ, which was enhanced for OBT. In contrast, the negative coherence_*ps*_, in the LPZ during PV turned positive during OBT in the patient groups, i.e. glaucoma and RP, while it remained negative, albeit reduced, for the controls. This reductions might be associated with some heterogeneity evident with the controls; specifically, for two controls LPZ responses turned positive for OBT compared to PV (see suppl. Figure S2; C6 & C7). We repeated the experiment on one of these participants a second time and reproduced the effect.

**Fig 4.**
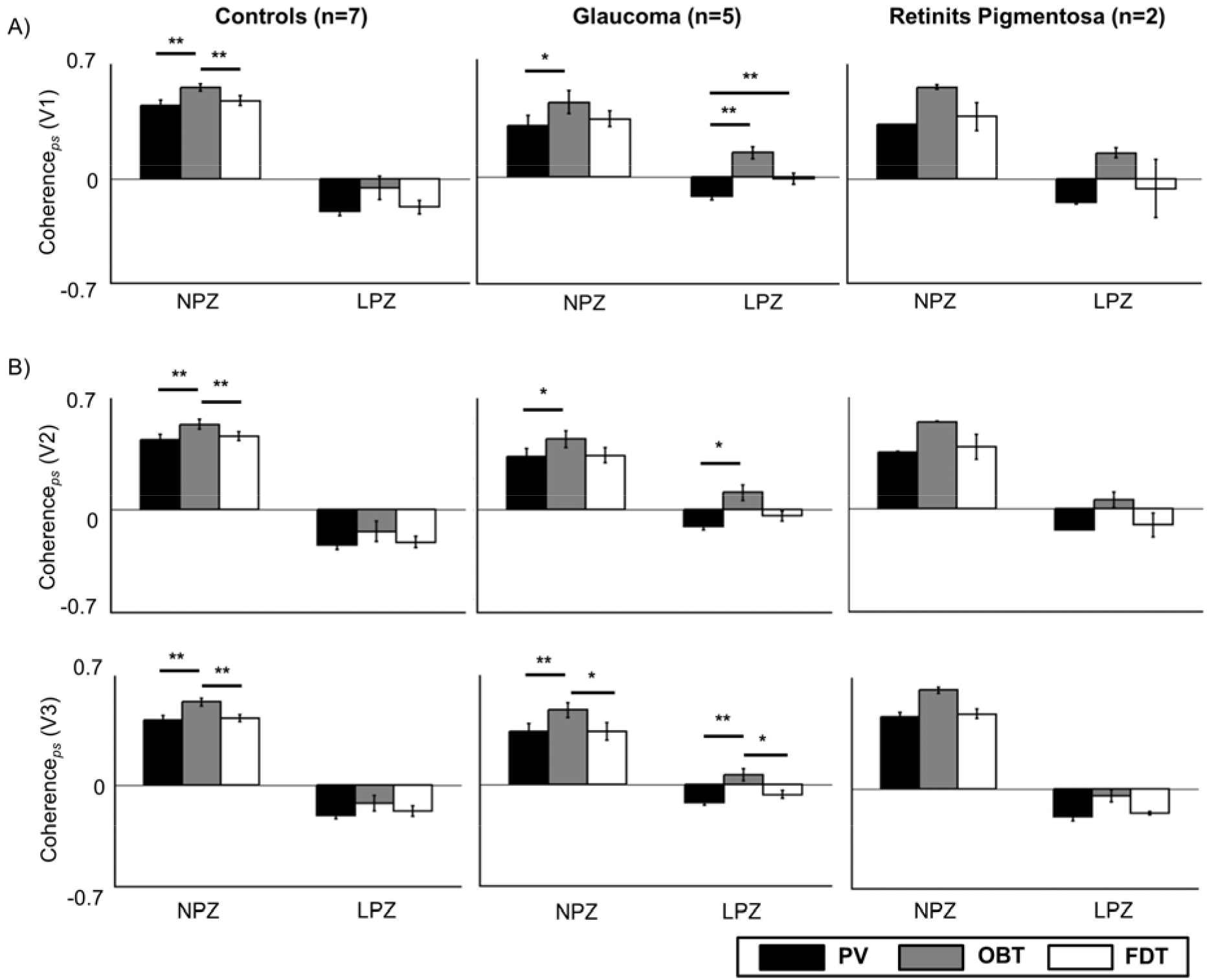
Task dependence of coherence_*ps*_ in V1 (A), V2 (B), and V3 (c) for the control (n=7), glaucoma (n=5), and retinitis pigmentosa groups (n=2; mean ± SEM). Controls and patients differ specifically for LPZ. Only in the patient groups’ LPZ, negative coherence_*ps*_ during PV are shifted to positive values for OBT. Significance test (corrected paired t-tests) on task effects were performed for glaucoma and controls, * p<0.05 & ** p<0.01. PV – passive viewing; OBT – one-back task; FDT – fixation-dot task.

For the control and the glaucoma groups the significance of these effects was determined with 2-way repeated measures ANOVAs (Factor 1 *task* and Factor 2 *ROI*), which specifically tested the coherence_*ps*_ difference between (i) PV and OBT, (ii) PV and FDT, and (iii) the equivalence of OBT and FDT as detailed in Table 2 (for post-hoc tests see Figure 4). (i) PV and OBT. *Task* and *ROI* were significant for both the control and the glaucoma group, while the interaction *task* x *ROI* was significant only for the latter. Post-hoc paired t-tests (corrected), further revealed enhanced responses during OBT in NPZ for both control [t = 4.4; p = 0.01] and glaucoma [t = 3.9; p = 0.03], but in LPZ only for glaucoma [t = 6.4; p = 0.006]. The main effect of ROI is anticipated as the two ROIs (NPZ & LPZ) differ in their coherence_*ps*_ to the stimulus. The task effect in the controls is driven by the change in amplitude of only NPZ responses, but by both NPZ and LPZ task-related modulations in the glaucoma group. This dependence of task effect on the ROI explains the significant interaction (*task* x *ROI*) reported only in glaucoma. (ii) PV and FDT. *ROI* was significant for both groups, *task* and the interaction *task* x *ROI* only for the glaucoma group. Post-hoc tests further revealed enhanced responses during FDT only for LPZ in glaucoma [t = 6.2; p = 0.006]. However, FDT did not evoke a net task-related shift from negative to positive coherence_*ps*_ in the LPZ. With respect to the initial rationale, i.e. whether PV is distinct from OBT, even in the absence of an alternative task as for FDT, PV appears to be sufficiently distinct. (iii) OBT and FDT. *ROI* and *task* were significant for both groups, the interaction *task* x *ROI* was marginally significant only for the glaucoma group. Post-hoc tests further revealed reduced responses during FDT only for NPZ in controls [t = 4.4; p = 0.01]. Similar to PV and OBT comparison, the significant difference in control NPZ in this case is likely a result of task-related amplitude change in the fMRI responses. The marginally significant interaction and non-significant LPZ differences in the glaucoma group indicates that FDT is less effective than PV in achieving passive viewing with respect to the stimulus.

**Table 2.** Two-way Repeated measures ANOVA of the reported coherence_*ps*_ measures for visual area V1. Factor 1: *Task* (Passive viewing, One-back task, Fixation-dot task); Factor 2: *ROI* (NPZ, LPZ). Three independent ANOVAs were done with different task combinations.

| Group | Main effect |  |  |  | Interaction |  |
| --- | --- | --- | --- | --- | --- | --- |
|  | Task |  | ROI |  | F | p |
|  | F | p | F | p |  |  |
| ANOVA 1 - Factor 1: *Task* (Passive viewing, One-back task); Factor 2: *ROI* (LPZ, NPZ) |  |  |  |  |  |  |
| Glaucoma | 28.9 | 0.006 | 31.6 | 0.005 | 37.3 | 0.004 |
| Controls | 15.8 | 0.007 | 163.8 | <0.001 | 0.58 | 0.480 |
| ANOVA 2 - Factor 1: *Task* (Passive viewing, Fixation-dot task); Factor 2: *ROI* (LPZ, NPZ) |  |  |  |  |  |  |
| Glaucoma | 24.5 | 0.008 | 33.5 | 0.004 | 15.5 | 0.017 |
| Controls | 1.3 | 0.292 | 192.4 | <0.001 | 0.05 | 0.832 |
| ANOVA 3 - Factor 1: *Task* (One-back task, Fixation-dot task); Factor 2: *ROI* (LPZ, NPZ) |  |  |  |  |  |  |
| Glaucoma | 9.2 | 0.039 | 30.3 | 0.005 | 8.5 | 0.044 |
| Controls | 7.1 | 0.037 | 166.7 | <0.001 | 0.89 | 0.383 |

As we had only two RP patients, we did not perform further statistics on their data. Nonetheless, we found task-dependent activity in the peripheral de-afferented visual cortex identical to the glaucoma cohort (Figure 4A). This is consistent with the results from (Masuda et al., 2010) and adds evidence to the replicability potential of such responses. For the FDT, one patient had positive coherence to the stimulus in the LPZ and the other negative (suppl Figure. S2).

Assessing whether the above task-dependence in V1 also applies to V2 and V3 we observed similarly significant differences between PV and OBT in the extra-striate LPZ representations in glaucoma patients, i.e. a shift from negative to positive responses (Figure 4B & see suppl. Table S1 for statistics). We further checked in these patients for any differential extent of task-dependence in V1, V2 and V3 by calculating the effect size (computed as Cohen’s d) of the observed task-related coherence_*ps*_. modulations in the LPZ. We observed a stronger effect size in V1 (d = 0.73) and relatively lower effect size for V2 (d = 0.55) and V3 (d = 0.51).

## 4. Discussion

We report that activations in the LPZ of V1, V2 and V3 were strongly related to performing a one-back task in glaucoma and retinitis pigmentosa (RP), but not in controls with simulated LPZ. This indicates that the limited remapping, previously reported for V1 in RP and macular degeneration (MD), is also a feature of V1 to V3 in glaucoma. These results thus suggest that strong limits of bottom-up plasticity are a general feature of the early human visual cortex that was de-afferented due to acquired lesions at the level of the retinal photoreceptors or ganglion cells.

While the response pattern we observed, i.e. LPZ activation for visual stimulus-locked tasks, has up-to-date not been reported for glaucoma, it fits well into the context of other studies on visual plasticity in acquired retinal defects [for e.g. MD (Baseler et al., 2011; Masuda et al., 2008), RP (Masuda et al., 2010)]. From the initial apparent heterogeneity of reports on the scope of plasticity in the LPZ of human V1, ranging from the absence of relevant cortical remapping (Baseler et al., 2011, Sunness et al., 2004) to large-scale reorganization (Baker et al., 2008; Dilks et al., 2009), the picture of limited bottom-up plasticity in human V1 has eventually emerged (Morland, 2015; Wandell and Smirnakis, 2009). Accordingly, Masuda et al. observed LPZ-responses in V1 during the performance of stimulus-related visual tasks and concluded that they were associated with task-dependent feedback instead of bottom-up plasticity in MD (Masuda et al., 2008) and RP (Masuda et al., 2010). In fact, the findings from our study replicate the findings by Masuda et al. for the rare disease RP and extend them to the much more prevalent disease, glaucoma.

### 4.1 Early visual cortex stability and plasticity

The potential of remapping in V1 in acquired defects is often inferred from adult animal models (Giannikopoulos and Eysel, 2006; Gilbert and Wiesel, 1992; Kaas et al., 1990). While some degree of developmental V1 plasticity has been reported for congenital vision disorders (Baseler et al., 2002), the nature and magnitude of a large-scale reorganization in adulthood is questionable (Wandell and Smirnakis, 2009) and still warrants quantitative ascertainment. The differential comprehension in the above literature on LPZ activation primarily arises from the variable definitions of cortical reorganization and remapping (Morland, 2015). While the proponents of plasticity speculate that the mere presence of abnormal LPZ responses is sufficient evidence, the critiques point out the need for such responses to be non-explainable by the normal visual cortex organization following visual field loss. In the context of the latter definition of cortical reorganization, from our data, bottom-up large-scale reorganization appears an unlikely cause of the reported LPZ activation in glaucoma, as it would lead to LPZ responses irrespective of the condition, i.e. task. While this supports top-down effects as a cause of the task-related LPZ responses, these do not appear to strictly follow the inverse visual hierarchy, i.e. a decreasing effect size from V3 to V1, as might be expected for top-down modulations. In fact, the differential activation (reflected by Cohen’s d) we observed in V1 was not exceeded by those in V2 and V3. A stronger effect size in the extra-striate areas (i.e. V3>V2>V1) would have added further evidence to a top-down hypothesis. Consequently, further research is necessary to decipher the nature of the top-down modulations. One rewarding avenue to pursue for this purpose is paved by the advent of MRI at submillimeter resolution (Fracasso et al., 2018; Kashyap et al., 2018; Ress et al., 2007; Yakupov et al., 2017). It opens the possibility to recover information on the directionality of information flow in the cortex via laminar imaging (Dumoulin et al., 2018; Lawrence et al., 2019) that allows to dissociate activations in cortical input and output layers. Consequently, future studies measuring layer-specific functional activity in the visual cortex might unravel the missing pieces of information, i.e. origin and directionality of task-related LPZ activations, to validate or invalidate existing theories on the aberrant cortical activity observed in patients with de-afferented visual cortex.

### 4.2 Clinical relevance in the context of emerging therapeutic interventions

A subset of glaucoma patients continue advancing towards blindness regardless of disease management. While emerging restoration therapies might offer treatment options for this patient entity, it has been suggested that vision loss associated changes at the level of visual cortex might be a reason for treatment failure in such cases (Davis et al., 2016; Gupta and Yücel, 2007; Nuzzi et al., 2018). Importantly, the existence of cortical responses in the de-afferented visual cortex, as demonstrated in the present study, suggests that the LPZ is still to some degree operational. This finding is consequently in support of approaches restoring visual input to the cortex (Aguirre, 2017; Beauchamp et al., 2020; Chuang et al., 2014; Jutley et al., 2017; Sena and Lindsley, 2017; Venugopalan et al., 2016). High cost and limited availability makes the utility of MRI in routine clinical ophthalmological examinations less plausible. However, our findings indicate the relevance of investigating the functional processing of the visual cortex in post-retinal lesions in preparation of demanding vision restoration procedures, e.g. to predict their clinical effectiveness.

### 4.3 Limitations and future directions

As a limitation in the study we acknowledge the small sample size of glaucoma patients, which was still sufficient to assess the relevant effects both at the individual and at the group level. It should be noted, that our target patients were those with strongly advanced VF deficits. Consequently, as glaucoma is an age-progressive disorder, such patients are mostly in their later stages of life and likely to have at least one MRI-related contraindication, which makes them a rare cohort for fMRI investigations. Future research should investigate the dynamics of the observed task-dependent LPZ responses in patients with different stages of the pathology and more importantly aim to uncover the mechanisms underlying such responses with submillimeter laminar fMRI imaging. This research should also include controls, as we observed in a minority of controls task-dependent effects for the simulated LPZ that resembled those found in the patients. At this point of time the underlying framework for this observation can only be speculated on, e.g. it might be related to aging or subclinical degenerations. Consequently, it disserves further assessment in future research.

## 5. Conclusion

In summary, we demonstrated in patients with advanced glaucoma, the existence of aberrant cortical responses in the supposedly de-afferented regions of the early visual cortex. The fMRI modulations are more likely to be driven by task-elicited top-down neural mechanisms than bottom-up cortical reorganization. Given similar findings in RP and MD, the results are indicative of a general mechanism behind such aberrant cortical responses that is not specific to the pathophysiology of the disease. We believe that these insights are of importance for the development of treatment and rehabilitation schemes in glaucoma and beyond.

## Acknowledgements

This project was supported by European Union’s Horizon 2020 research and innovation programme under the Marie Sklodowska-Curie grant agreements No.675033 (EGRET plus) and by the German research foundation (DFG: HO2002/20-1) to MBH. The funding organization did not have any role in the study design, collection, analysis and interpretation of the data, or publication of this research. We also thank Katharina Jürse for her help with the recruitment of participants and Denise Scheermann for her help with scanning.

## CRediT authorship contribution statement

Gokulraj T. Prabhakaran: Conceptualization, Methodology, Formal analysis, Investigation and Data curation, Writing-original draft. Claus Tempelmann: Methodology, Writing - review and editing. Markus Wagner: Investigation, Writing-review and editing. Hagen Thieme: Writing - review and editing. Michael B. Hoffmann: Conceptualization, Methodology, Supervision, Writing - review and editing, Funding acquisition. Khaldoon O. Al-Nosairy: Investigation, Writing - review and editing.

