## Supplementary material for "Functional dynamics of de-afferented early visual cortex in glaucoma": suppl. Table S1

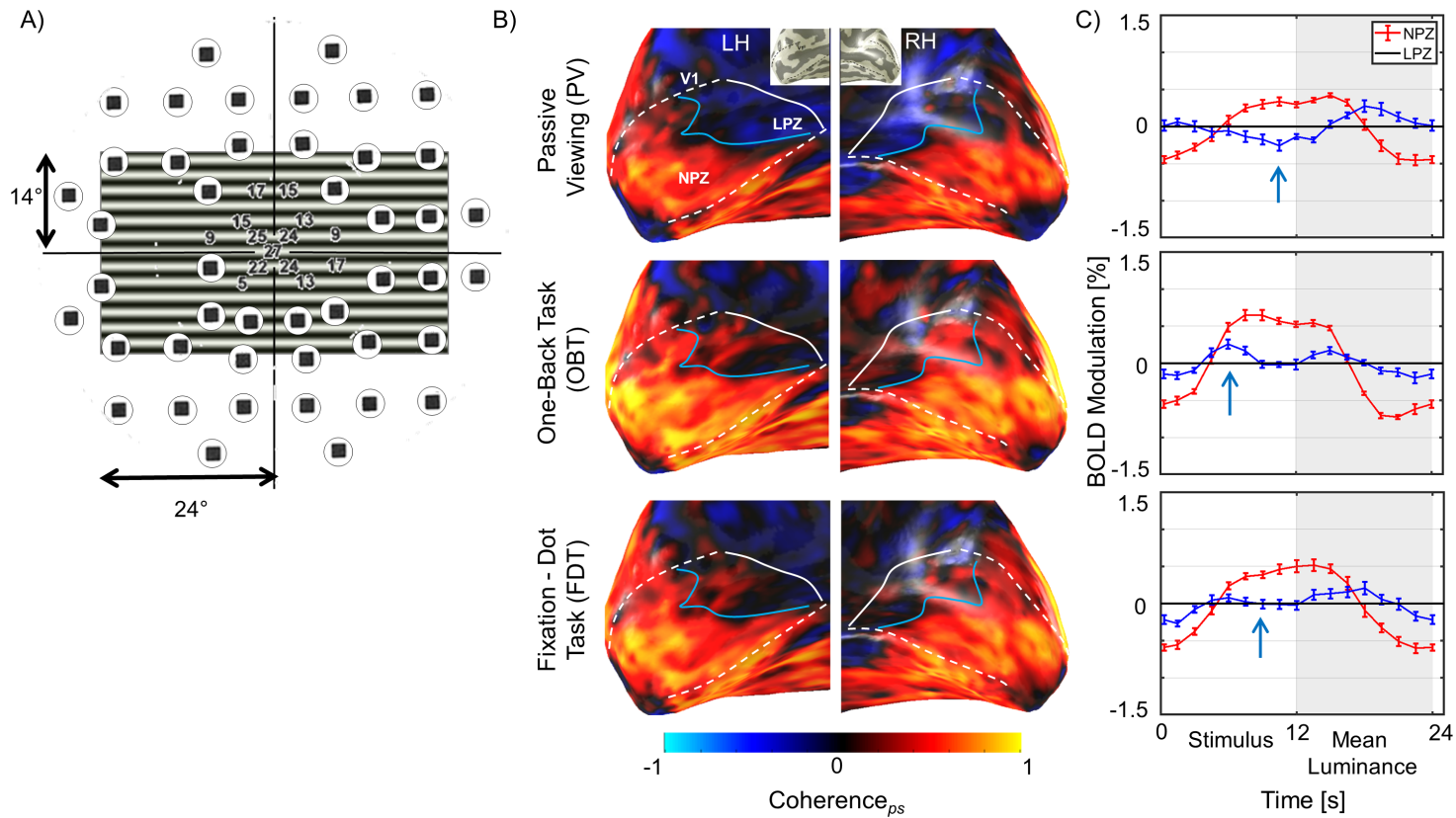

**Fig S1. fMRI-activations for RP2.** **(A)** Visual field in the left eye (stimulated in the fMRI experiment) as determined perimetrically superimposed approximately on stimulus layout illustrates the peripheral visual field defect in this patient. **(B)** BOLD-activations (coherence*_ps_*) as false-color overlays onto the inflated occipital lobe. **(C)** Average single-cycle BOLD time series. Depending on tasks, LPZ responses are negative (PV) or positive (OBT) or reduced (FDT), as indicated by arrows. Conventions as for Figure 2.

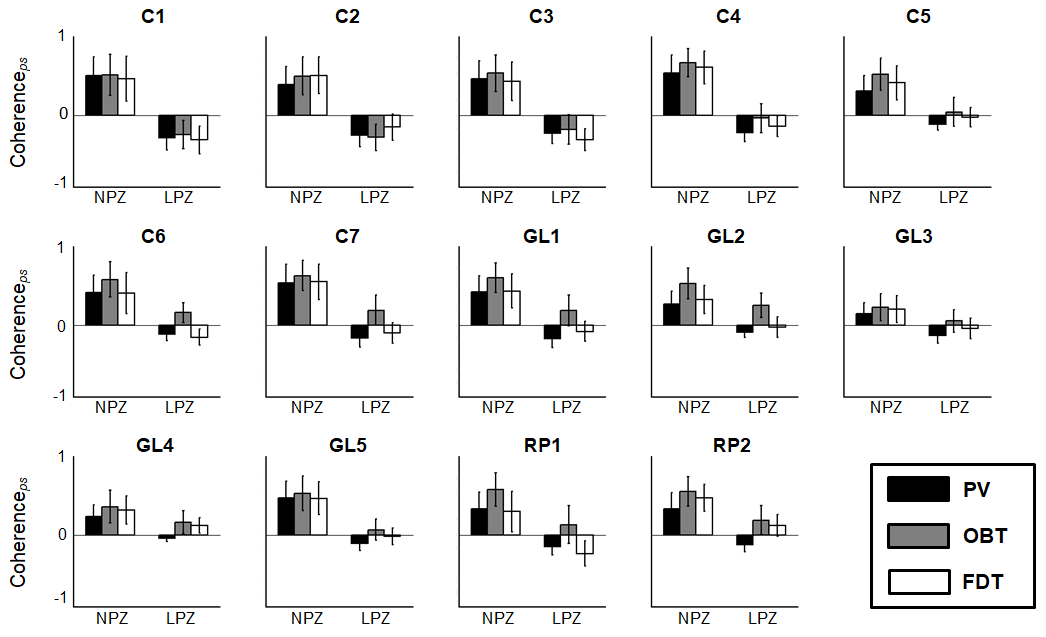

**Fig S2. Individual task dependences of coherence*_ps_* in V1 (mean±SD).** Interindividual variations of the task dependence are particularly evident for controls, where C6 and C7 show task dependence in LPZ. Conventions as for Figure 4.

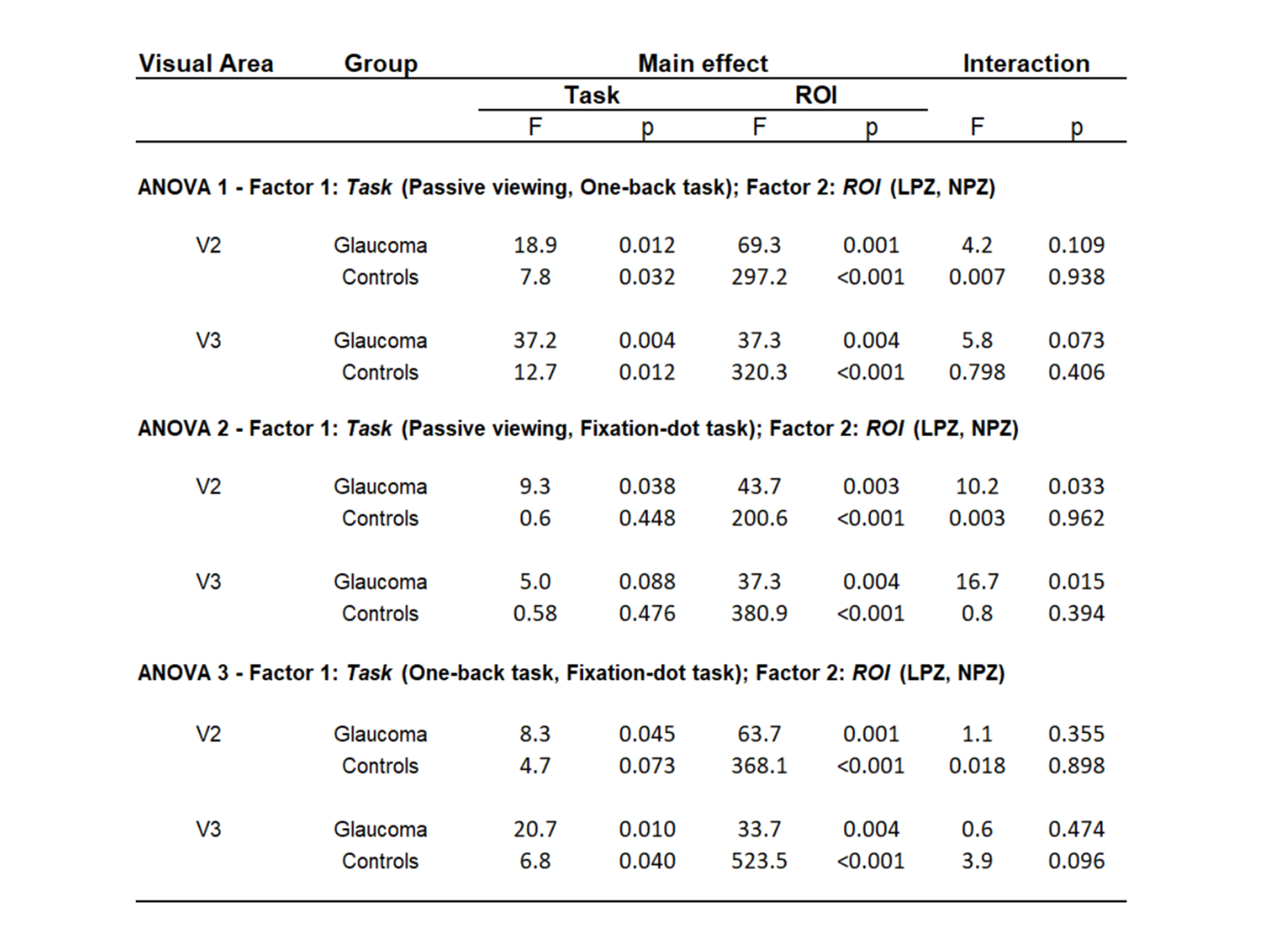

**Supp Table S1. Two-way repeated measures ANOVA** of the reported Coherence*_ps_* measures for visual area V2 & V3. Factor 1: *Task* (Passive viewing, One-back task, Fixation-dot task); Factor 2: *ROI* (NPZ, LPZ). Three independent ANOVAs were done with different task combinations.
